## Supplementary Figures and Tables for "Agonistic CD40 antibody therapy induces tertiary lymphoid structures but impairs the response to immune checkpoint blockade in glioma"

#### SUPPLEMENTARY MATERIAL

### Extended Data Figure legends

**Extended Data Fig. 1: Induction of tertiary lymphoid structures (TLS) in murine glioma models. (a)** Immunofluorescent staining of dense $CD45^+B220^+$ TLS adjacent to the meninges and in close proximity to the tumor tissue (dense nuclei, bottom) in GL261 tumor-bearing mice. Scale bar: 200 µm. **(b)** Schematic representation of TLS location in murine glioma models. TLS typically formed in the tumor-bearing hemisphere and were observed in multiple locations in close contact with the pia mater around the cortex or in the choroid plexuses. **(c-d)** $CD45^+B220^+$ TLS were not present in the brain of (**c**) non-treated healthy mice or (**d**) mice which had undergone mock tumor implantation followed by treatment with αCD40 antibodies. Scale bars: 200 µm. **(e-g)** Immunostaining of TLS in the brain of αCD40-treated GL261 tumor-bearing mice revealed the presence of (**e**) rare $GFP^+CD11c^+$ dendritic cells, (**f**) $Foxp3^+$ T regs and (**g**) $F4/80^+$ macrophages. For clarity, the same area within the dotted square in (e) is shown on the bottom right of the panel in the absence of B220 and CD3 channels. The arrow indicates a $CD11c^+$ dendritic cell positive for GFP (or "tumor", white). The magenta arrows in (f) indicate cells that stained triple-positive for CD3, CD4 and FoxP3. Red arrows indicate cells that stained positive for CD3 only. Scale bars: 50µm. **(h-i)** Quantification of (**h**) the number of TLS per section and (**i**) TLS surface area plotted against the time of sacrifice for αCD40-treated GL261 tumor-bearing mice. **(j)** Representative images of dense $CD45^+B220^+$ clusters before (top panel) and after (bottom panel) laser capture microdissection. Scale bars: 50 µm. **(k)** Gene expression of *Ccl21* in laser capture micro-dissected $CD45^+B220^+$ clusters, compared with laser capture micro-dissected tumor tissue and normal brain tissue. $n_{(\text{of LMD areas})}$=4-6/group. ANOVA with Tukey's multiple comparison correction. Values: mean. **(l-o)** Immunofluorescent stainings of αCD40-induced TLS in the CT-2A model showing TLS composition and organization. Arrows in (l) indicate a majority of $Ki67^+$ T cells (white arrows) and one $Ki67^+$ B cell (yellow arrow). Scale bars: 50 µm.

**Extended Data Fig. 2: B cells were necessary for αCD40-mediated TLS induction. (a)** Immunofluorescent staining showing cells in the tumor area which are positive for the therapeutic αCD40 antibody. **(b)** Representative images showing the difference in composition between a TLS and a T cell aggregate in the meninges. TLS showed a clear B cell core surrounded by fewer T cells and scattered DCs, while T cell aggregates were infiltrated with only few B cells and comprised of a clear network of DCs. Scale bars: 50 µm. **(c)** Quantification of aggregates of T cells close to the meningeal layer in the indicated treatment groups. n=5-8 mice/group. ANOVA with Tukey multiple correction. Values: mean.

**Extended Data Fig. 3: TLS composition and location in glioma patients. (a-d)** Immunohistochemical staining showing the presence of rare (**a,c**) $CD23^+$ follicular B cells and (**b,d**) $CD138^+$ plasma cells in immature and organized TLS. Zoom areas in (a-d) are indicated by black squares and the relative zoom panels are shown to the right of each image. Scale bars: 50µm. **(e)** A dense cluster of nuclei *in the depth of a cerebral sulcus, identified* in a section obtained from a WHO Grade II oligodendroglioma (patient 6, Supplementary Table 1) that stained positive for mutated isocitrate dehydrogenase 1 (IDH1). *The b*lack square area in (e) is magnified to the right. Arrow in (e): cerebral sulcus. Arrow in zoom panel: *dense cluster of nuclei.* Scale bar: 5 mm. **(f)** Schematic representation of the locations were TLS were observed in human glioma samples: 1) in close proximity to the meninges, 2) in the white matter, close to the tumor and 3) inside the tumor tissue. For each location, a representative immunohistochemical staining is shown. This includes an overview of the tissue, where TLS location is indicated by a black square. A zoom of the square area is shown to the right. Scale bars: 2 mm.

**Extended Data Fig. 4: T cell characterization in αCD40-treated mice. (a)** Representative images of T cell infiltration in αCD40-treated GL261 tumors that stained positive or negative for TLS. Scale bar: 50 µm. **(b)** Quantification of $CD25^+FoxP3^+$ Tregs as a percentage of $CD45^+$ cells in GL261 tumors treated with rIgG2a or αCD40. n=8mice/group. **(c)** Representative FACS plots of the $CD3^+CD4^+CD25^+FoxP3^+$ T cell population quantified in (b).

**Extended Data Fig. 5: Systemic response to αCD40 therapy.** Panels **(a-f)** show quantifications obtained from spleens of GL261 and CT-2A tumor-bearing mice treated with rIgG2a or αCD40 antibodies. **(a,d)** Quantification of $CD62L^+CD44^-$ T cells as a percentage of $CD8^+$ cells. **(b,e)** Quantification of $CD62L^-CD44^+$ T cells as a percentage of $CD8^+$ cells. **(c,f)** Quantification of $CD127^-KLRG1^+$ cells as a percentage of $CD8^+CD44^+$ T cells. **(g-q)** Systemic cytokine levels of (**g**) TNFα, (**h**) IFNγ, (**i**) CXCL10, (**j**) IL-1β, (**k**) IL-2, (**l**) IL-6, (**m**) IL-12p70, (**n**) IL-16, (**o**) IL-23 and (**p**) IL-5 in the serum of GL261 tumor-bearing mice on day 13, day 19 and day 25 post tumor implantation, after treatment with rIgG2a or αCD40 antibodies. The dashed line indicates the lower limit of detection. Statistics were performed with *t*-test or multiple t-tests with Sidak-Bonferroni's correction for multiple comparisons. *p<0.05, **p< 0.01, ***p < 0.001, ****p < 0.0001. Values: mean. (a-f) n=5-7mice/group). (g-p) n=6mice/group.

**Extended Data Fig. 6: αCD40 induced a dysfunctional T cell phenotype in glioma-bearing mice. (a)** Example of how generations of proliferating $CD8^+$ T splenocytes were defined based on cell trace violet staining. Each peak represents one generation. Generation 0 corresponds to the highest intensity peak, where cells did not divide. Generation 6 corresponds to the lowest intensity peak, where cells have divided roughly 6 times. **(b)** Schematic illustration of the experimental layout used to obtained data shown in panels (c-d). In brief, GL261 glioma-bearing mice were treated with rIgG2a or αCD40 antibodies on days 10, 13, 16 and 19 post-tumor implantation. On day 22, the mice were sacrificed and $CD8^+$ T cells were isolated from the brain, stained with cell trace violet (CTV) to assess proliferation and re-stimulated *in vitro* with concanavalin A (ConA) for 24h and 72h. (**c**) Histogram showing the proportion of $CD8^+$ T cells in each proliferation peak based on CTV staining. (**d**) Histogram showing CD69 protein levels on the $CD8^+$ T cell population. **(e)** Schematic illustration of the experimental layout used to obtained data shown in panels (f-h). In brief, GL261 glioma-bearing mice were treated with rIgG2a or αCD40 antibodies on days 10, 13, 16 and 19 post-tumor implantation. On day 22, the mice were sacrificed and $CD45^+$ T cells were isolated from the brain and co-cultured with GL261 cells in vitro for 24 and 72h. **(f-g)** Percentages of (**f**) $CD107a^+$ and (**g**) $IFN\gamma^+$ $CD8^+$ T cells in rIgG2a and αCD40 groups at the indicated time-points. n=2mice/group. **(h)** Relative viability of GL261 cells after 72h of co-culture. n=2mice/ group.

**Extended Data Fig. 7: αCD40 impairs the response to PD-1 and CTLA-4 checkpoint blockade.** All data shown in this figure was obtained from GL261 glioma-bearing mice. **(a)** Representative image of TLS in the indicated treatment groups. Scale bar: 50µm. **(b)** Quantification of the $CD45^+$ surface area of each individual TLS for the indicated treatment groups. n=8-17mice/group. **(c)** Serum levels of rat antibodies (Rat IgG) in the indicated treatment groups on day 19 post tumor implantation. n=4mice/group. ANOVA with Tukey's multiple correction. **p<0.01. Values: mean. **(d)** Kaplan–Meier survival curve of mice treated with a single dose of αCD40 on day 9 or treated with a single dose of αCD40 on day 9 followed by three doses of αPD1 antibodies on days 10, day 13 and day 16 (as indicated by red and blue arrows, respectively). n=9-10mice/group. Log rank test; *p < 0.05, ***p<0,001. **(e)** Quantification of the number of TLS for the indicated treatment groups. n=6-9mice/group. **(f)** Kaplan–Meier survival curve of mice treated with αCTLA-4 antibodies and/or αCD40 antibodies on days 10, day 13, day 16 and day 19 (as indicated by arrows). n=16/group. Log rank test; **p < 0.01. **(g)** Representative image of TLS in the indicated treatment groups. Scale bar: 50µm. **(h,i)** Quantification of (**h**) the number and (**i**) the $CD45^+$ surface area of each individual TLS for the indicated treatment groups. n=4-9mice/group.

**Extended Data Fig. 8: αCD40 activated brain-infiltrating dendritic cells and myeloid cells.** Phenotypic analysis of myeloid subtypes in GL261 glioma-bearing mice. **(a,b)** Quantification of (**a**) $CD19^-CD11b^+CD11c^+$ dendritic cells (DCs) and (**b**) $CD19^-CD11b^+CD11c^-$ myeloid cells as a percentage of immune cells in the brain of mice in the indicated treatment groups. n=4-7mice/group. **(c)** Heatmap showing the expression levels of activation and immunosuppression markers on DCs and myeloid cells in the brain of mice in the indicated treatment groups.

n=4-7mice/group. **(d-e)** Quantification of IL-12$^+$ cells as a percentage of (**d**) DCs and (**e**) myeloid cells in the brain of mice in the indicated treatment groups. n=4-7mice/group. **(f-g)** Quantification of IL-10$^+$ cells as a percentage of (**f**) DCs and (**g**) myeloid cells in the brain of mice in the indicated treatment groups. n=4-7mice/group. Statistics was performed with ANOVA with Tukey's multiple correction. *p<0.05, **p<0.01, ****p<0,0001. Values: mean.

**Extended Data Fig. 9: αCD40 therapy did not increase the expression of soluble immunosuppressive factors in B cells.** Analysis of regulatory molecule on B cells in GL261 tumors. **(a)** CD5$^+$CD1d$^+$ positive cells as a percentage of CD19$^+$B220$^+$ B cells on day 20 and on day 25 post tumor implantation, with or without 5h stimulation with PMA and Ionomycin (P/I). n=7-8mice/group. *t*-test, **p<0.01, ***p<0.001. **(b)** Representative plots of the CD1d$^+$CD5$^+$ B cell population quantified in (a). **(c-f)** Gene expression of *Il10, Tgfb*1, *Ccl22* and *Lgals1* on B cells sorted on day 20 post tumor implantation. n=8mice/group. *t*-test, *p<0.05, ****p<0.0001. **(g-j)** Gene expression of *Il10, Tgfb*1, *Ccl22* and *Lgals1* on B cells sorted on day 25 post tumor implantation. n=8mice/group. *t*-test.

**Extended Data Fig. 10: αCD40 therapy induced CD11b$^+$ B cells and suppressed CD4$^+$ T cells.** All figure panels show data from GL261 glioma-bearing mice. **(a)** Quantification of CD11b$^+$ cells as a percentage of B cells in the spleen, in the indicated treatment groups. n=4mice/group. **(b)** Mean fluorescence intensity (MFI) of CD11b on CD11b$^+$ myeloid cells versus CD11b$^+$ B cells in the brain. Data were merged from rIgG2a, αCD40, αPD1 and αCD40+αPD1 treatment groups. n=23 mice. (**c**) Systemic levels of IL-10 on day 13, day 19 and day 25 post tumor implantation, after treatment with rIgG2a or αCD40 antibodies, with or without B cell depletion using an αCD20 antibody. The dashed line indicates the lower limit of detection. The data point marked as an empty red square was determined to be an outlier using the Grubbs test, thus it was excluded from the statistical analysis. n=6mice/group. Statistics was performed with multiple *t*-test. **(d)** Quantification of CD69$^+$ cells as a percentage of CD4$^+$ T cells in the brain, in the indicated treatment groups. n=5-8mice/group. **(e)** Mean fluorescence intensity (MFI) of Ki67 in CD4$^+$ T cells in the brain, in the indicated treatment groups. n=5-8mice/group. Statistics were performed with either *t*-test or ANOVA with Tukey's multiple correction if not indicated otherwise. *p<0.05, **p<0.01, ****p<0.0001. Values: mean.

**Extended Data Fig. 11: CD11b expression on B cells inhibits CD8$^+$ T cell responses. (a)** Schematic illustration of the experimental layout used to obtained data shown in panels (b-g). In brief, B cells were isolated from wt C57BL/6 mice and co-cultured with LPS for 48h to induce upregulation of CD11b. LPS-treated B cells were then co-cultured for 72h with splenocytes in the presence of a control antibody (rIgG2b k) or a CD11b neutralizing antibody. Data on B and T cells were collected by FACS. **(b)** Mean fluorescence intensity (MFI) of CD11b on B cells with or without LPS stimulation. **(c-f)** Quantification of (**c**) CD69$^+$, (**d**) CD107a$^+$ (**e**) IFNγ$^+$ and (**f**) proliferating cells as a percentage of CD8$^+$ T cells after co-culture with LPS-induced B cells in the presence of rIgG2b k or an αCD11b antibody. **(g)** Representative histogram showing the shift of proliferation peaks on CD8$^+$ T cells co-cultured with B cells in the presence of a control antibody (rIgG2b k) or of a CD11b-neutralizing antibody. n=3mice/group. *t*-test; *p<0.05, ****p<0.0001.

**Extended Data Fig. 12: Representative example of gating strategy for flow cytometry analysis.**
The panel shows the gating strategy for flow cytometry analysis of CD45$^+$ cells isolated from the brain of glioma-bearing mice and stained for T cell markers. **(a)** Out of the total events recorded, debris were excluded from the starting population based on FSC-A vs SSC-A values. **(b)** Doublets were excluded from the population shown in (a) based on SSC-W values. **(c)** Cells that stained positive for the Live-Dead staining were excluded from the population shown in (b) to select for live cells. **(d)** Out of all live cells, CD45$^{high}$ cells were gated on to select for the immune cell population. **(e)** T cells were identified by gating for CD3$^+$ cells on the CD45$^{high}$ population. **(f)** Cytotoxic T cells were identified by gating for CD8$^+$CD4$^-$ cells on the CD3$^+$ population. **(g)** Effector cytotoxic T cells were identified by selecting CD44$^+$ cells out of the CD8$^+$CD4$^-$ population shown in (e). Gate is based on FMO

control (gray: FMO; blue: sample). **(h)** The CD44$^{+}$ population shown in (g) was divided in four subpopulations based on the expression levels of CD127 and KLRG1. Gate based on FMO controls.

#### Extended Data Figures

**EXTENDED DATA FIGURE 1**

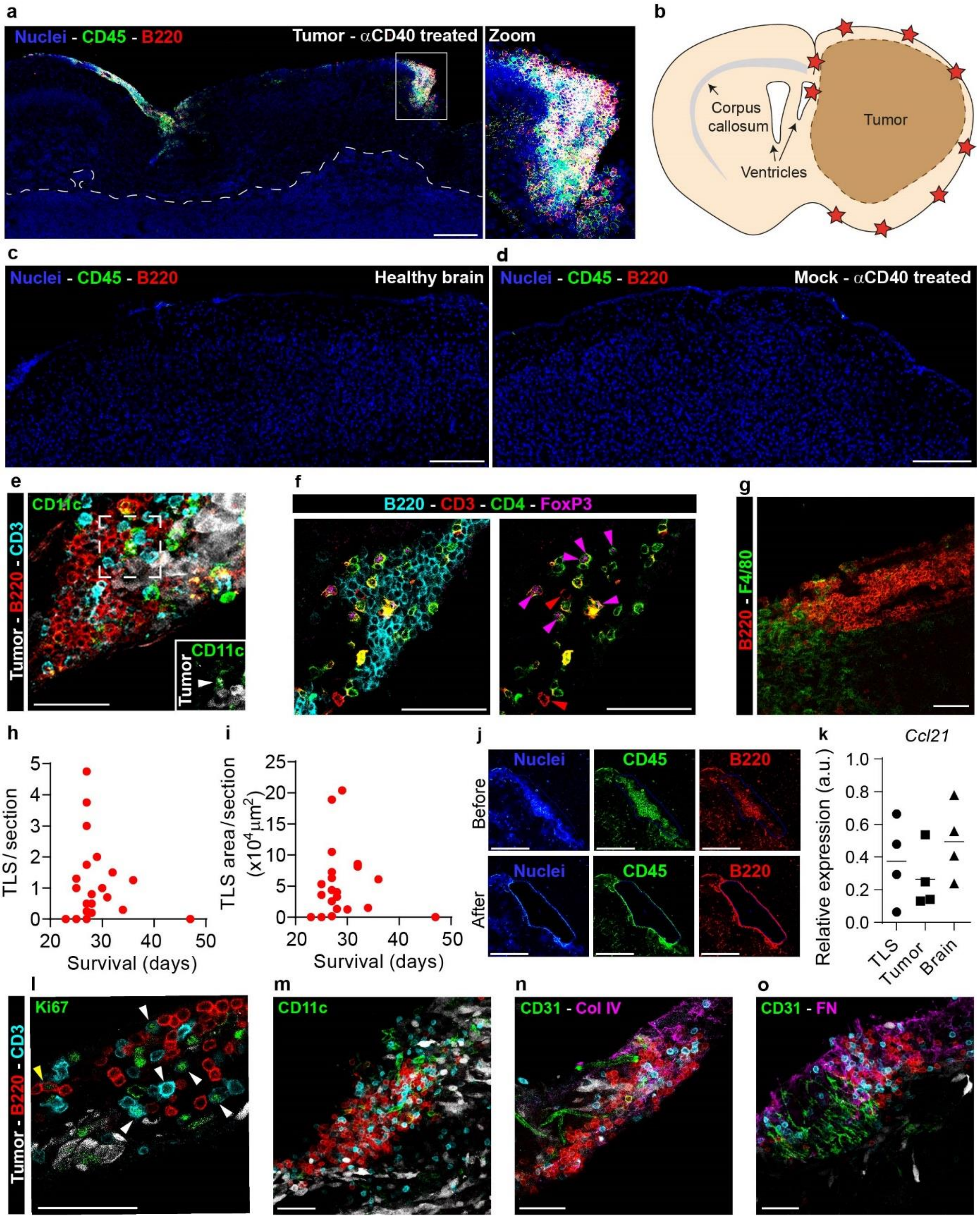

**EXTENDED DATA FIGURE 2**

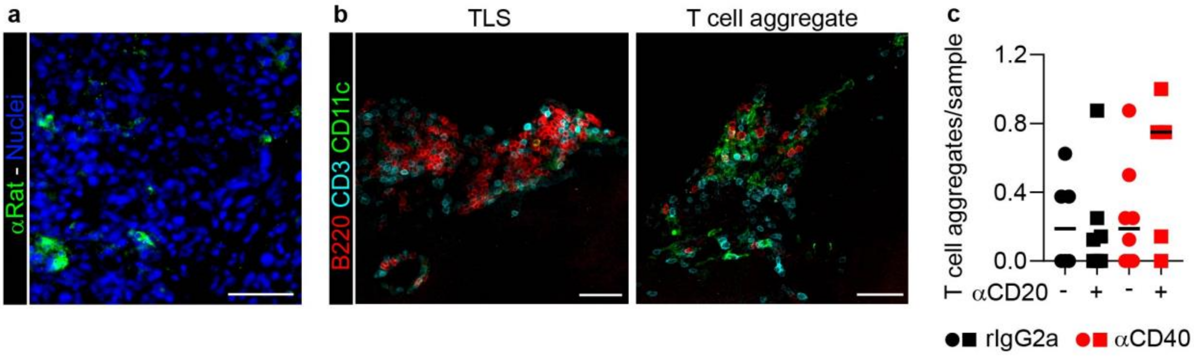

### EXTENDED DATA FIGURE 3

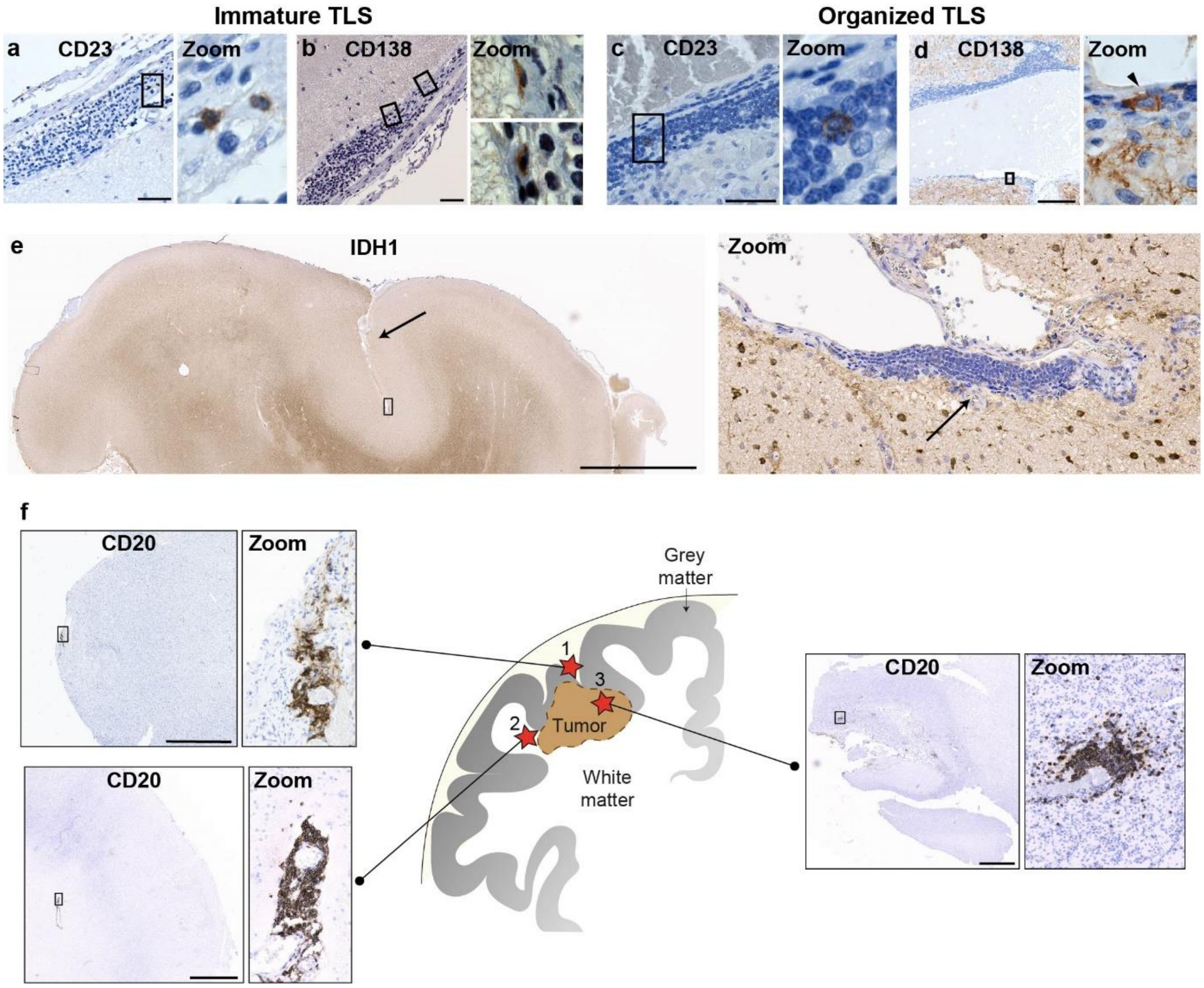

**EXTENDED DATA FIGURE 4**

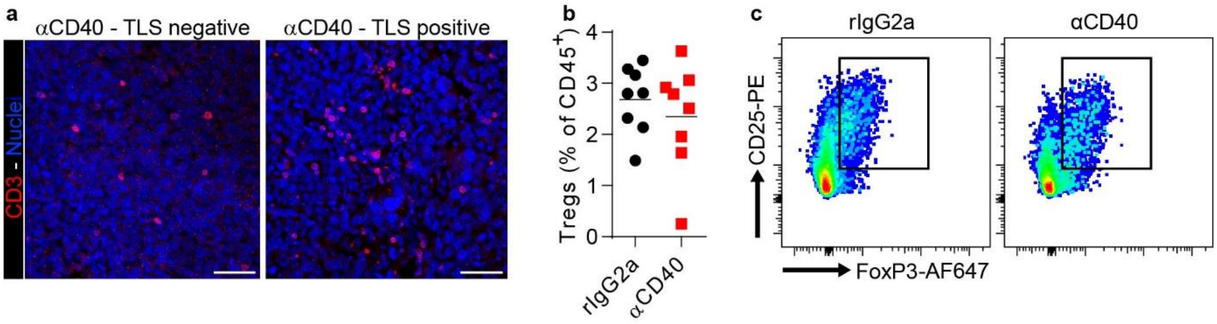

**EXTENDED DATA FIGURE 5**

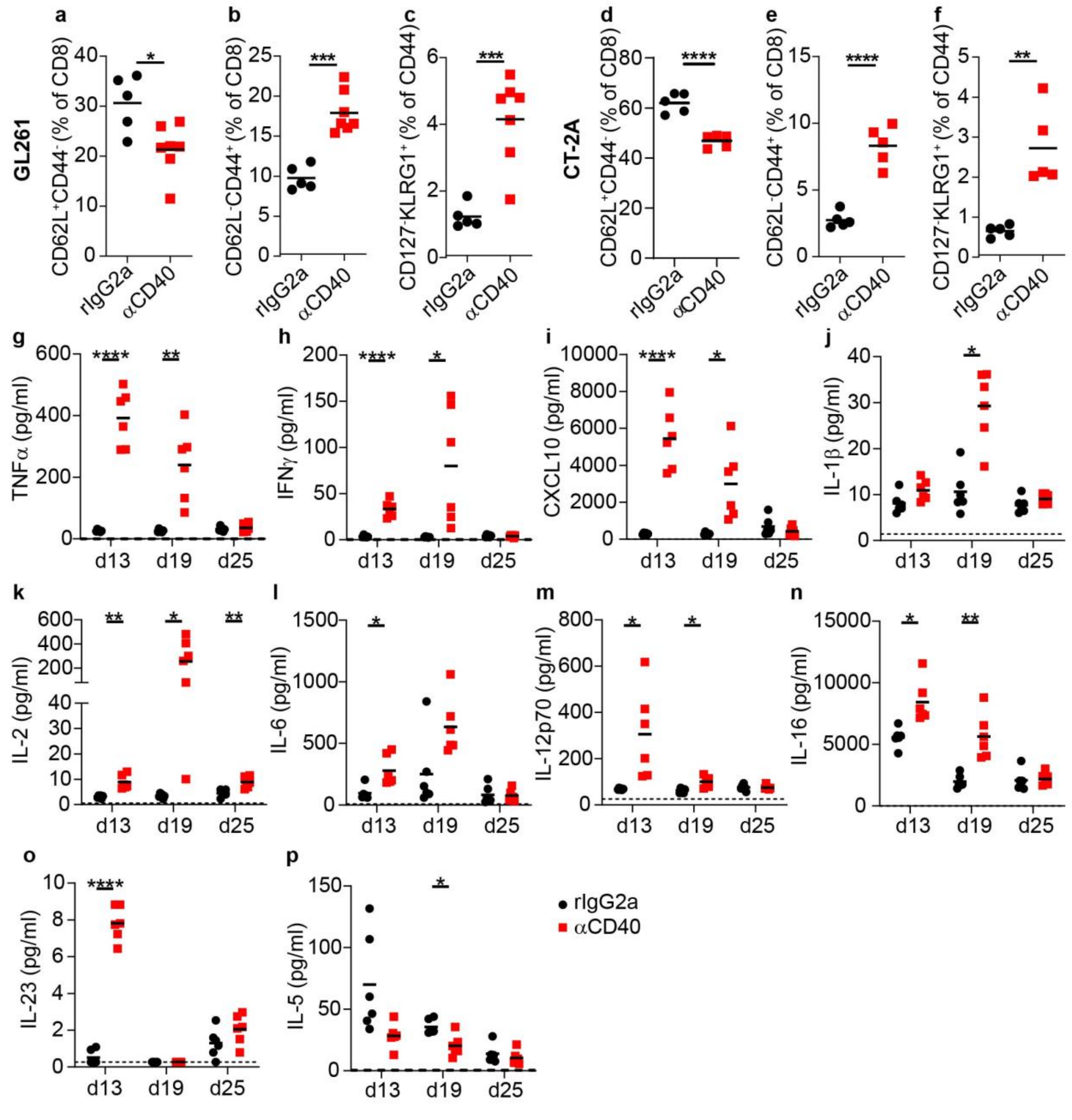

**EXTENDED DATA FIGURE 6**

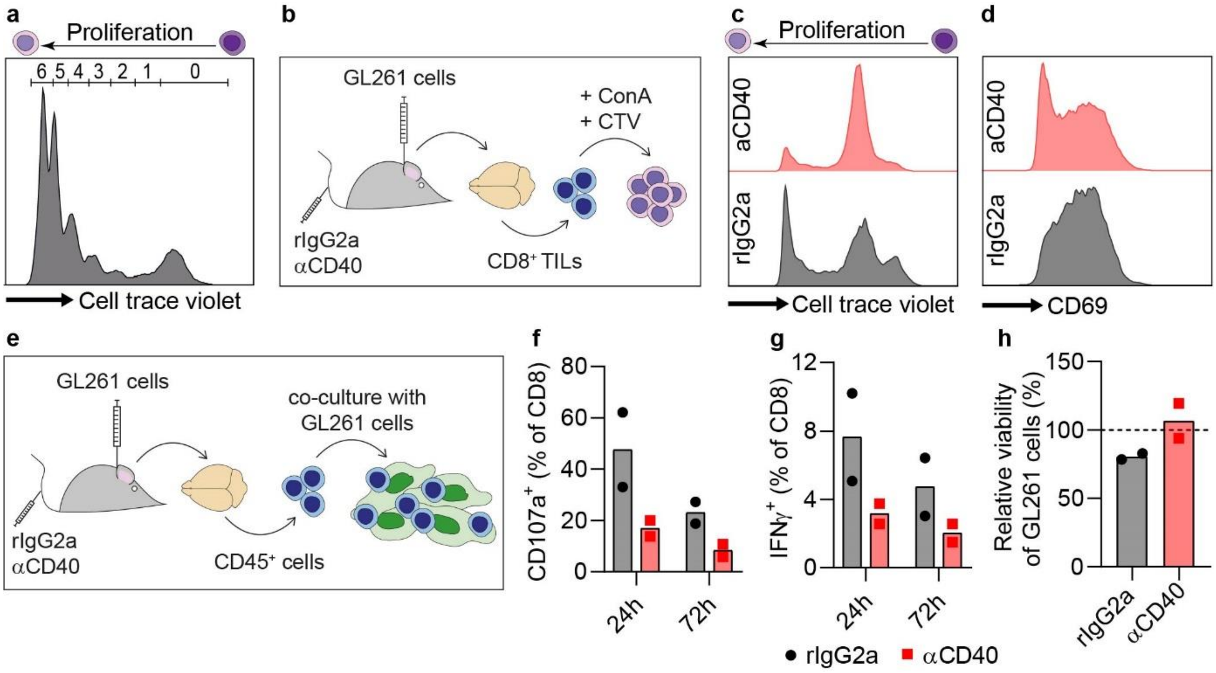

**EXTENDED DATA FIGURE 7**

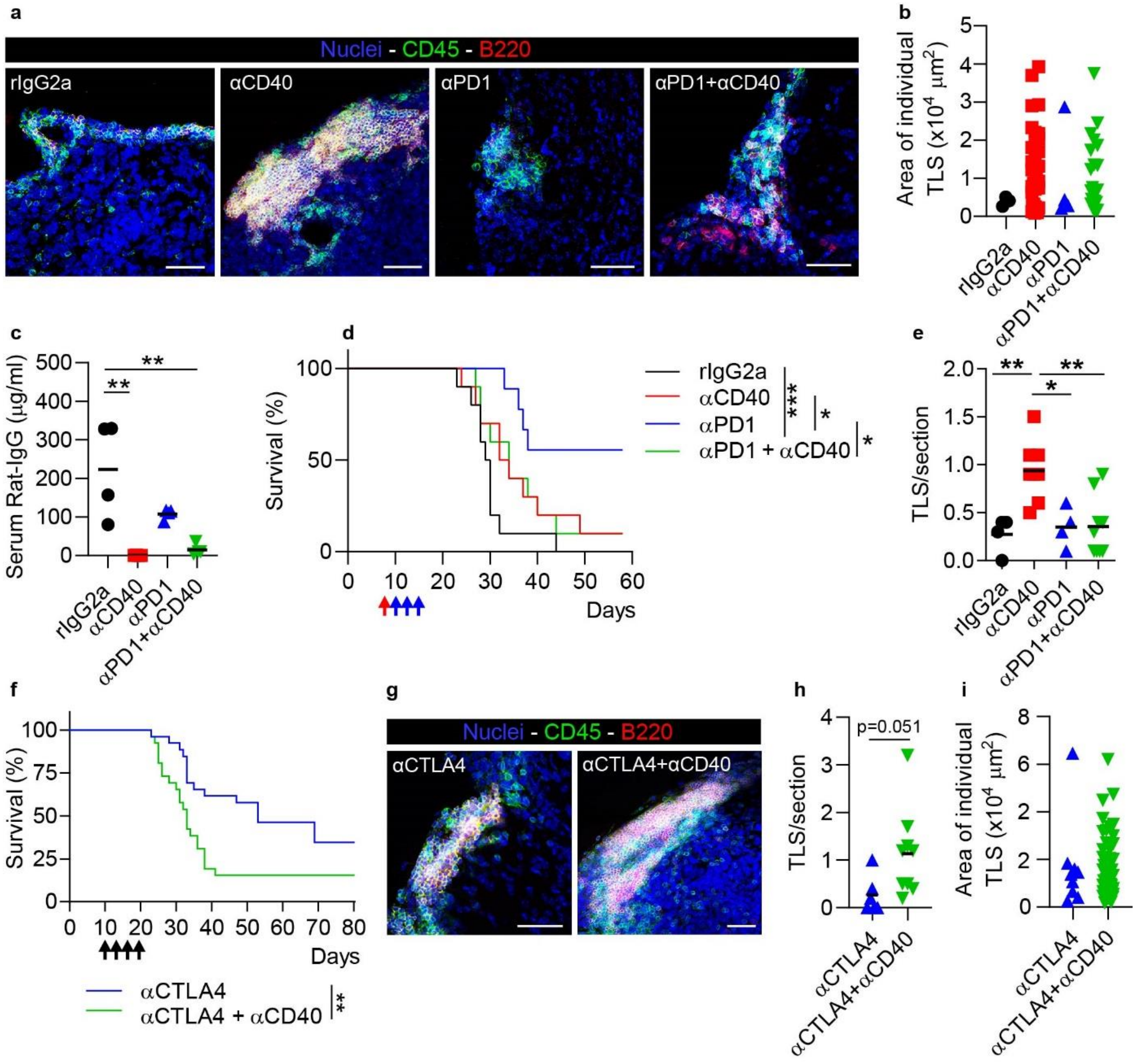

**EXTENDED DATA FIGURE 8**

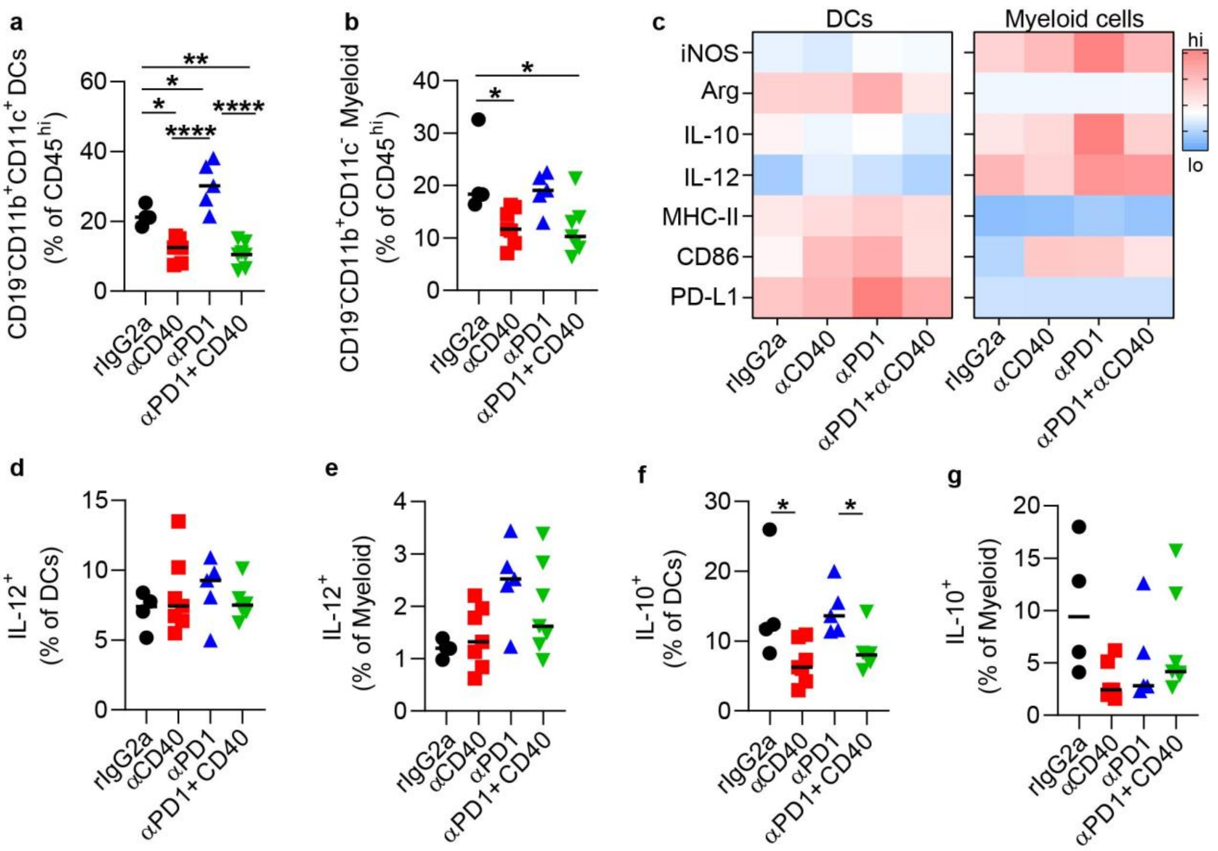

**EXTENDED DATA FIGURE 9**

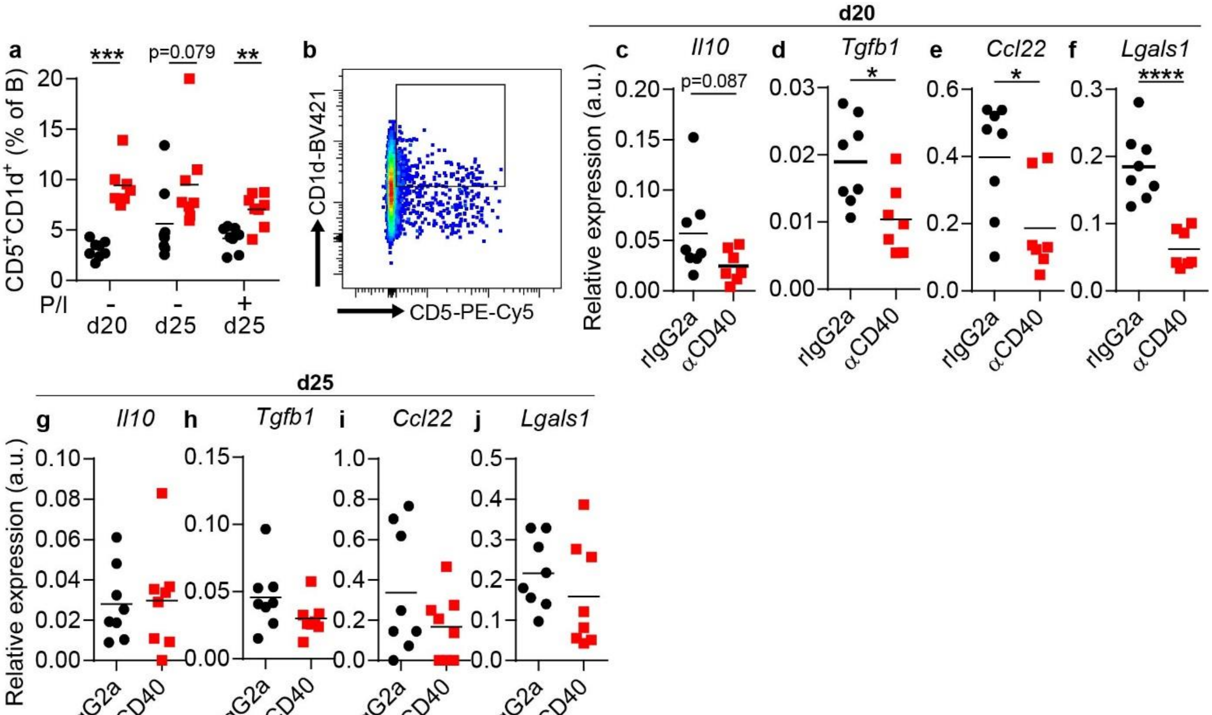

### EXTENDED DATA FIGURE 10

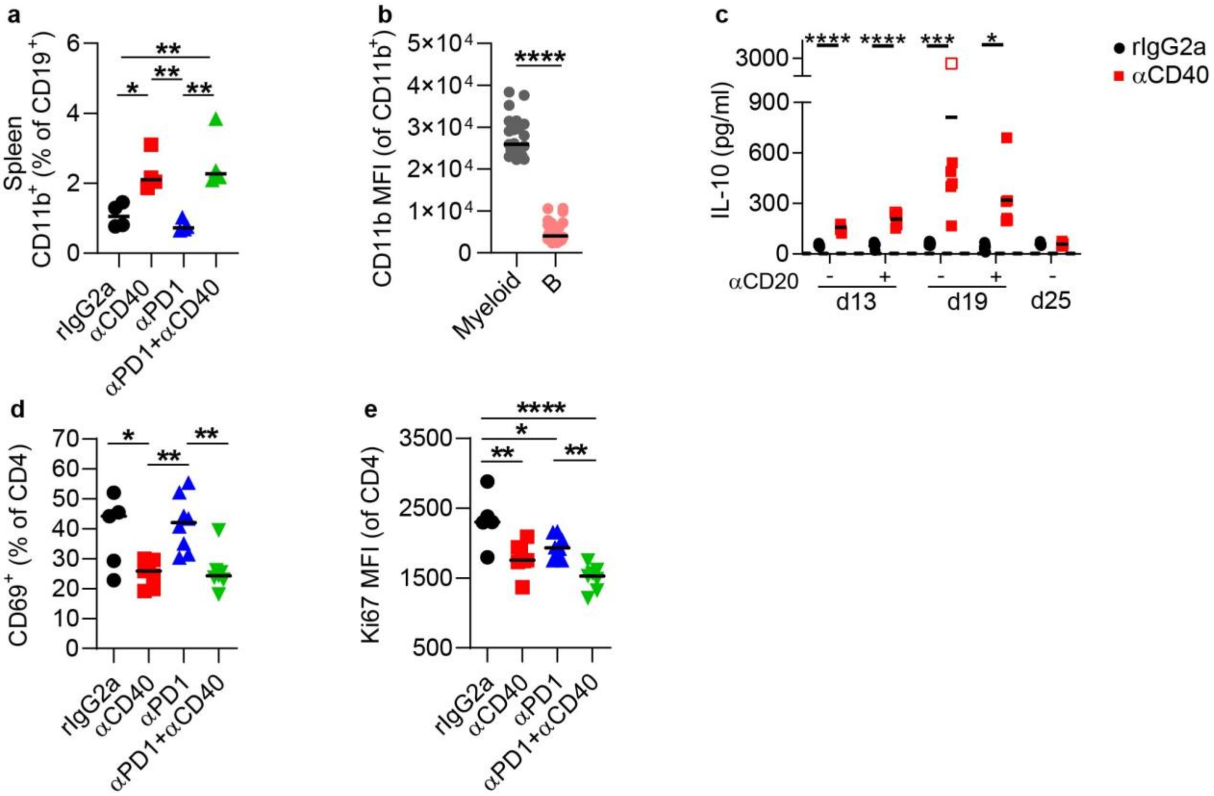

**EXTENDED DATA FIGURE 11**

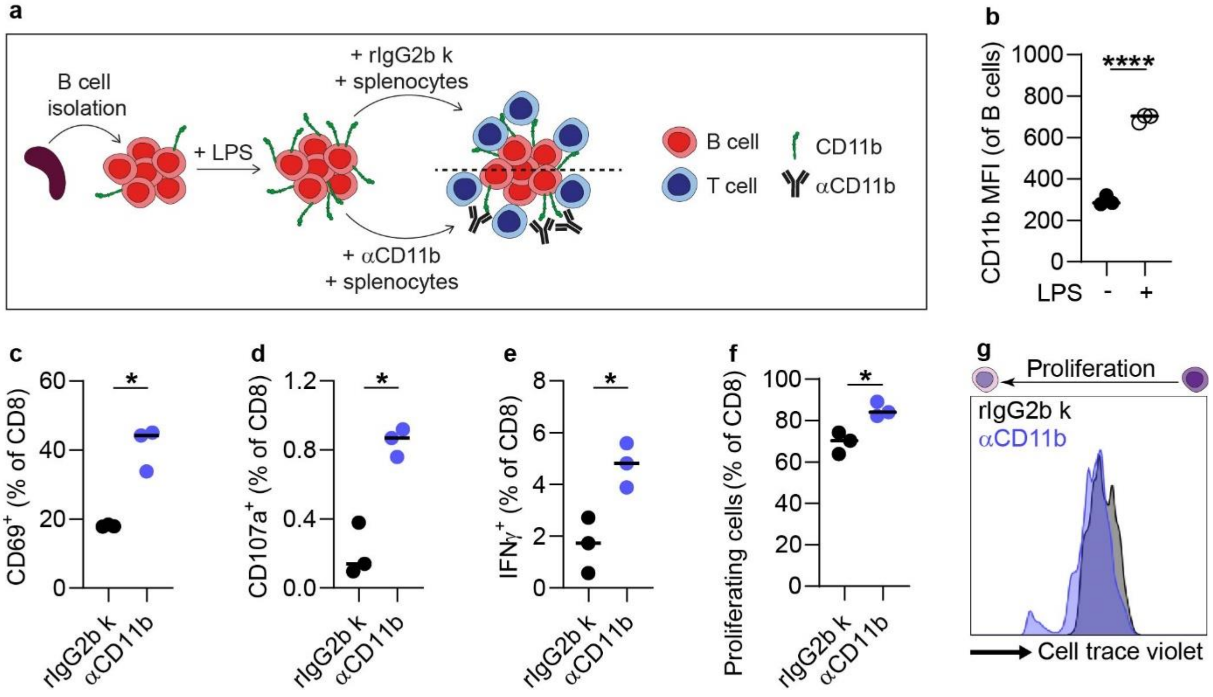

**EXTENDED DATA FIGURE 12**

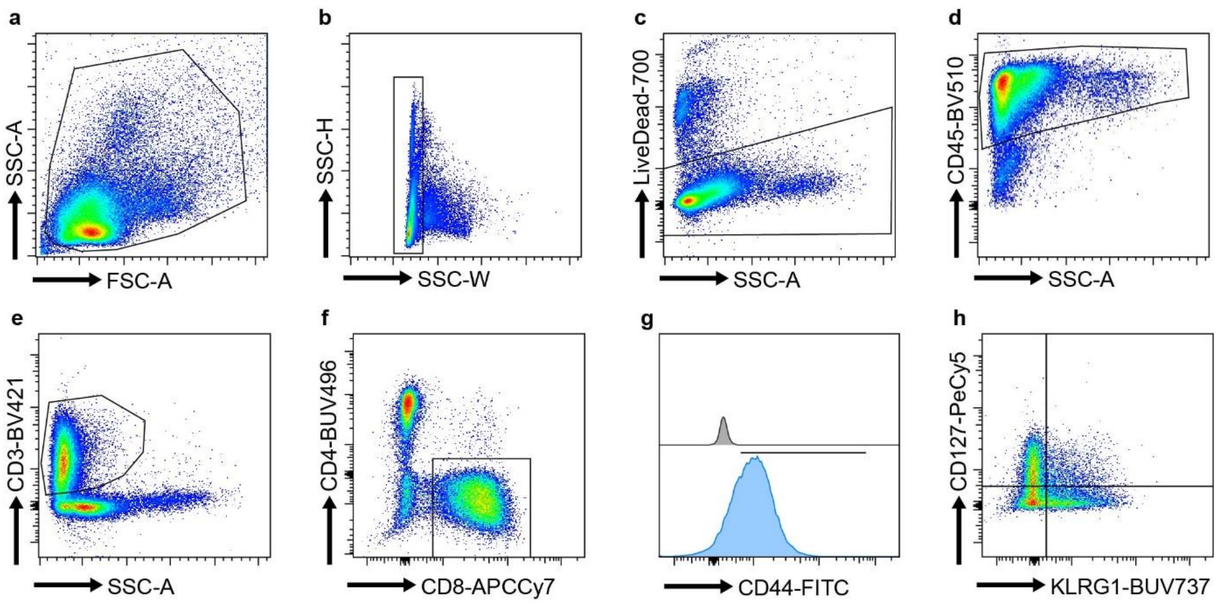

#### Supplementary Tables

**Supplementary Table 1.** Histological characteristics and WHO classification of the human glioma samples included in the cohort. The table also reports TLS presence and classification for each patient. Abbreviations: mut (mutation); co-del (co-deletion); NA (not assessed).

| Patient ID | IDH1 mut | 1p/19q co-del | ATRX mut | p53 mut | Histological diagnosis (WHO) | Grade | Type of section | TLS presence | Immature TLS | Organized TLS |
|---|---|---|---|---|---|---|---|---|---|---|
| **1** | - | - | - | - | Astrocytoma | II | En block, large | - | 0 | 0 |
| **2** | + | - | + | - | Astrocytoma | II | En block, large | - | 0 | 0 |
| **3** | + | - | + | - | Astrocytoma | II | En block, large | - | 0 | 0 |
| **4** | + | + | - | - | Oligodendroglioma | II | En block, large | - | 0 | 0 |
| **5** | + | + | - | - | Oligodendroglioma | II | En block, large | + | 0 | 3 |
| **6** | + | + | - | - | Oligodendroglioma | II | En block, large | + | 3 | 0 |
| **7** | - | - | - | - | Anaplastic astrocytoma | III | En block, large | - | 0 | 0 |
| **8** | + | + | - | - | Anaplastic oligodendroglioma | III | En block, large | + | 6 | 1 |
| **9** | + | + | - | - | Anaplastic oligodendroglioma | III | En block, large | - | 0 | 0 |
| **10** | + | + | - | - | Anaplastic oligodendroglioma | III | En block, large | - | 0 | 0 |
| **11#** | - | - | - | - | Glioblastoma | IV | En block, large | + | 6 | 0 |
| **12** | - | - | NA | - | Glioblastoma | IV | Large biopsy | - | 0 | 0 |
| **13** | - | - | NA | + | Glioblastoma | IV | Large biopsy | + | 1 | 4 |
| **14** | - | - | NA | - | Glioblastoma | IV | Large biopsy | - | 0 | 0 |
| **15** | - | - | NA | + | Glioblastoma | IV | Large biopsy | - | 0 | 0 |
| **16** | - | - | NA | + | Glioblastoma | IV | Large biopsy | + | 1 | 3 |
| **17** | - | - | NA | + | Glioblastoma | IV | Large biopsy | - | 0 | 0 |
| **18** | - | - | NA | + | Glioblastoma | IV | Large biopsy | - | 0 | 0 |
| **19** | - | - | NA | + | Glioblastoma | IV | Large biopsy | - | 0 | 0 |
| **20** | - | - | NA | + | Glioblastoma | IV | Large biopsy | - | 0 | 0 |
| **21** | - | - | NA | - | Glioblastoma | IV | Large biopsy | + | 1 | 0 |
| **22** | - | - | NA | - | Glioblastoma | IV | Large biopsy | + | 1 | 0 |
| **23** | - | - | NA | - | Glioblastoma | IV | Large biopsy | + | 0 | 1 |
| **24** | - | - | NA | - | Glioblastoma | IV | Large biopsy | + | 2 | 0 |
| **25*** | - | - | - | - | Glioblastoma | IV | Large biopsy | + | 0 | 4 |
| **26** | - | - | - | - | Glioblastoma | IV | Large biopsy | - | 0 | 0 |

#Patient 11: this patient was also afflicted with chronic lymphocytic leukemia

*Patient 25: samples collected from a primary and a secondary surgery were available. TLS were found in both cases.

**Supplementary Table 2.** Mouse primer sequences used for real-time quantitative PCR

| Target | | Primer sequence (5' - 3') |
|---|---|---|
| *Cxcl13* | Fw | CATAGATCGGATTCAAGTTACGCC |
| | Rs | TCTTGGTCCAGATCACAACTTCA |
| *Ccl19* | Fw | ACCACACTAAGGGGCTATCAG |
| | Rs | TTCTTCAGTCTTCGGATGATGC |
| *Ccl21* | Fw | GCTGCAAGAGAACTGAACAGACA |
| | Rs | CGTGAACCACCCAGCTTGA |
| *Ccl22* | Fw | AGGTCCCTATGGTGCCAATGT |
| | Rs | CGGCAGGATTTTGAGGTCCA |
| *Il10* | Fw | GCTCTTACTGACTGGCATGAG |
| | Rs | CGCAGCTCTAGGAGCATGTG |
| *Lgals1* | Fw | CAAGCTGCCAGACGGACAT |
| | Rs | AGGCCACGCACTTAATCTTGA |
| *Lta* | Fw | GCATCTTCTAAGCCCTGGGGG |
| | Rs | TGTCATGTGGAGGACCTGCTGTG |
| *Ltb* | Fw | GTTCAACAGCTGCCAAAGGG |
| | Rs | CATCCAAGCGCCTATGAGGT |
| *Tgfβ* | Fw | CTCCCGTGGCTTCTAGTGC |
| | Rs | GCCTTAGTTTGGACAGGATCTG |
| *Tnfsf14* | Fw | AGCAGCACATCTTACAGGAGC |
| | Rs | AGCTGCACTTTGGAGTACACA |

**Supplementary Table 3.** List of antibodies used for immunostaining of mouse vibratome sections, cryosections and tissue for laser microdissection (LMD).

| | Marker | Clone or Cat# | Conjugated | Fluorochrome | Company |
|---|---|---|---|---|---|
| **Used for LMD** | B220 | RA3-6B2 | Yes | PE | Biolegend |
| | CD45 | 30-F11 | Yes | AF488 | Biolegend |
| **Used for cryo- or vibratome sections** | Anti-rat | A-21208 | Yes | AF647 | Life Technologies |
| | CD11b | M1-70 | Yes | APC | Biolegend |
| | CD11c | N418 | No | - | Abcam |
| | CD19 | ab227019 | No | - | Abcam |
| | CD21 | SC0681 | No | - | Thermo Fisher |
| | CD23 | PA5-79242 | No | - | Thermo Fisher |
| | CD3 | 17A2 | Yes | BV421 | BD |
| | CD31 | 2H8 | No | - | Life Technologies |
| | CD35 | 8C12 | Yes | DyLight 350 | Novus Biologicals |
| | CD45 | 30-F11 | Yes | APC | Biolegend |
| | CD62L | ME-14 | Yes | AF647 | Biolegend |
| | ColIV | ab6586 | No | - | Abcam |
| | FN | ab2415 | No | - | Abcam |
| | FoxP3 | 150D | Yes | AF647 | Biolegend |
| | F4/80 | BM8 | Yes | PeCy5 | Biolegend |
| | Ki67 | ab16667 | No | - | Abcam |

**Supplementary Table 4.** List of antibodies and dyes used to characterize T cells and antigen presenting cells (APCs) in the tumor microenvironment (TME) by flow cytometry.

| | Marker | Clone | Conjugated | Fluorochrome | Company |
|---|---|---|---|---|---|
| **17-color panel used to characterize T cells** | CD45 | 30-F11 | Yes | BV510 | BD |
| | CD3e | 145-2C11 | Yes | BV421 | BD |
| | CD8 | 53-6.7 | Yes | APC-Cy7 | BD |
| | CD4 | GK1.5 | Yes | BUV496 | BD |
| | CD44 | IM7 | Yes | FITC | Biolegend |
| | CD62L | MEL-14 | Yes | PE-Cy7 | BD |
| | CD69 | H1.2F3 | Yes | BB700 | BD |
| | CD127 | A7R34 | Yes | PE-Cy5 | Biolegend |
| | PD-1 | 29F.1A12 | Yes | BV785 | Biolegend |
| | TIM-3 | RMT3-23 | Yes | BV605 | Biolegend |
| | LAG-3 | C9B7W | Yes | BV711 | BD |
| | Ki67 | 16A8 | Yes | PE dazzle594 | Biolegend |
| | CXCR5 | L138D7 | Yes | BV650 | Biolegend |
| | KLRG1 | 2F1 | Yes | BUV737 | BD |
| | Foxp3 | 150D | Yes | AF647 | Biolegend |
| | CD25 | 3C7 | Yes | PE | Biolegend |
| | LiveDead | - | - | Fixable viability dye 700 | BD |
| **17-color panel used to characterize APCs** | CD45 | 30-F11 | Yes | BV510 | BD |
| | IL10 | JES5-16E3 | Yes | PE dazzle594 | Biolegend |
| | CD19 | 6D5 | Yes | APC-Cy7 | Biolegend |
| | IA/IE | M5/114.15.2 | Yes | BB700 | BD |
| | CD11b | M1/70 | Yes | BUV395 | BD |
| | Ly6G | 1A8 | Yes | BV421 | BD |
| | CD11c | N418 | Yes | PE-Cy5 | Biolegend |
| | CX3CR1 | SA011F11 | Yes | BV650 | Biolegend |
| | Ly-6C | HK1.4 | Yes | BV785 | Biolegend |
| | CD103 | M290 | Yes | BV711 | Biolegend |
| | F4/80 | 6F12 | Yes | BV605 | BD |
| | PD-L1 | 10F.9G2 | Yes | PE | Biolegend |
| | IL-12 (p40/p70) | C15.6 | Yes | APC | BD |
| | iNOS | CXNFT | Yes | A488 | eBioscience |
| | Arginase I | A1exF5 | Yes | PE-Cy7 | eBioscience |
| | CD86 | GL-1 | Yes | BUV737 | BD |
| | LiveDead | - | - | Fixable viability dye 700 | BD |

**Supplementary Table 5.** List of additional antibodies and dyes used for flow cytometry or FACS.

| | Marker | Clone | Conjugated | Fluorochrome | Company |
|---|---|---|---|---|---|
| **Other antibodies used for flow cytometry or FACS** | CD45 | 30-F11 | Yes | FITC | Biolegend |
| | B220 | RA3-6B2 | Yes | AF488 | Biolegend |
| | B220 | RA3-6B2 | Yes | PE | Biolegend |
| | B220 | RA3-6B2 | Yes | APCCy7 | Biolegend |
| | CD1d | 1B1 | Yes | BV421 | BD |
| | CD19 | 6D5 | Yes | APC | Biolegend |
| | CD19 | 1D3 | Yes | PerCPCy5.5 | BD |
| | CD4 | GK1.5 | Yes | PerCP | Biolegend |
| | CD4 | RM4-5 | Yes | AF488 | Biolegend |
| | CD5 | 53-7.3 | Yes | PE-Cy5 | Biolegend |
| | CD107a | 1D4B | Yes | PECy7 | Biolegend |
| | CD69 | H1.2F3 | Yes | PE | BD |
| | CD69 | H1.2F3 | Yes | APC | Biolegend |
| | CD11b | M1/70 | Yes | APC | Biolegend |
| | CD19 | 6D5 | Yes | FITC | Biolegend |
| | CD3 | 145-2C11 | Yes | PerCPCy5.5 | Invitrogen |
| | CD8 | 53-6.7 | Yes | APCCy7 | BD |
| | IFNγ | XMG1.2 | Yes | PE | BD |
| | LiveDead | - | - | Zombie Aqua | Biolegend |
